# *Ex vivo* therapeutic base and prime editing using chemically derived hepatic progenitors in a mouse model of tyrosinemia type 1

**DOI:** 10.1101/2020.09.14.297275

**Authors:** Yohan Kim, Jihyeon Yu, Sung-Ah Hong, Jeongyun Eom, Kiseok Jang, Seu-Na Lee, Jae-Sung Woo, Jaemin Jeong, Sangsu Bae, Dongho Choi

## Abstract

DNA base editors and prime editing technology capable of therapeutic base conversion enable *ex vivo* gene editing therapy for various genetic diseases. For such therapy, it is critical that the target cells survive well both outside the body and after transplantation. In this regard, chemically derived stem/progenitor cells are attracting attention as the most useful cell sources for clinical trials. Here, we generate chemically derived hepatic progenitors from the hereditary tyrosinemia type1 model mouse (HT1-mCdHs) and successfully correct the disease causing mutation using both adenosine base editors (ABEs) and prime editing tools. After transplantation into HT1 mice, ABE-corrected HT1-mCdHs repopulated the liver with fumarylacetoacetate hydrolase-positive cells and dramatically increased the survival rate of HT1 model mice, suggesting a safe and effective *ex vivo* gene editing therapy.

## Introduction

Hereditary tyrosinemia type I (HT1), an autosomal recessive disorder caused by a deficiency in fumarylacetoacetase (FAH), results in liver failure due to the accumulation of toxic metabolites from the tyrosine metabolic pathway and can lead to hepatocellular carcinoma (HCC) (Chinsky et al., 2017; Nobili et al., 2010). Although 2-[2-nitro-4-trifluoromethylbenzoyl]-1,3-cyclohexanedione (NTBC) is used therapeutically, it does not treat the fundamental HT1 deficiency. Moreover, some patients lack NTBC sensitivity and a risk of HCC remains during therapy. Previously, several groups tried gene therapy approaches involving virus-mediated delivery of full-length *Fah* complementary DNA (cDNA) into the liver in HT1 (*Fah*^*-/-*^) model mice (Grompe et al., 1998; Overturf et al., 1997). However, these strategies have potential limitations: the exogenous gene, unaffected by the native chromatin structure of the endogenous locus, will be constitutively expressed, at levels that differ from that of the endogenous gene, and there is a possibility of insertional mutagenesis as a result of viral vector integration into the host genome.

In a different strategy, CRISPR-mediated therapeutic editing approaches have been applied to alter a second gene in the disease pathway, thereby lessening the effects of the *Fah* mutation. Pankowicz *et al*. showed that in HT1 model mice, Cas9 nuclease-mediated disruption of the hydroxyphenylpyruvate dioxygenase (*Hpd*) gene, which has a role in the second step of tyrosine catabolism, alleviated the disease symptoms such that the mice exhibited a benign HT3 phenotype. Employing a similar strategy, Rossidis *et al*. used a cytosine base editor, instead of Cas9, to disrupt the *Hpd* gene by inducing a premature termination codon in the middle of the gene. Both experiments involved the direct delivery of CRISPR-associated tools via lentivirus or adeno-associated virus (AAV), i.e., they represent *in vivo* gene editing strategies. Thus, their clinical application would involve potential challenges such as safety issues associated with viral delivery and the possibility of insertional mutagenesis caused by integration of the viral vectors. Recently, Song et al. demonstrated successful base conversion of the *Fah* gene mutation via hydrodynamic tail-vain injection of adenine base editors (ABEs), which mediate A to G conversion, with a non-viral delivery system. However, in this case the lack of an *in vivo* selection process to exclude cells containing CRISPR-mediated off-target effects may be a potential limitation to clinical applications.

Alternatively, *ex vivo* gene editing strategies can bypass the potential drawbacks of *in vivo* strategies; gene-corrected cells and cells validated to be free of off-target effects can be selected to increase efficiency and safety, respectively. Furthermore, recipients would not be exposed to CRISPR effectors, which may circumvent pre-existing immunological responses in humans(Crudele and Chamberlain, 2018). To date, a few studies have explored *ex vivo* gene editing strategies for applications in HT1 model mice, but most trials used primary hepatocytes (PHs) as the source cell for gene correction and transplantation (Hickey et al., 2016; VanLith et al., 2018). Because PHs cannot proliferate and maintain their function in an *in vitro* environment, after transplantation the corrected cells became engrafted with low efficiency and caused short-term gene rescue effects. In addition, lentiviral vectors or AAV were used for CRISPR delivery due to the low transfection efficiency in PHs, raising the possibility of virus-associated safety issues. Instead of PHs, differentiable cells such as embryonic stem cells (ESCs) (Basma et al., 2009; Rambhatla et al., 2003), induced pluripotent stem cells (Chen et al., 2012; Sullivan et al., 2010), mesenchymal stem cells (Banas et al., 2007; Lee et al., 2004), and direct converted cells (Huang et al., 2014; Sekiya and Suzuki, 2011) can alternatively be used as sources of cells for transplantation. However, those cells are associated with potential challenges for clinical applications, such as low differentiation efficiency, incomplete function of differentiated cells compared with primary cells (Song et al., 2009; Wu and Tao, 2012), immune rejection (Rong et al., 2014), risk of tumorigenesis (Lee et al., 2013; Miura et al., 2009), use of viral vectors (Huang et al., 2014; Yu et al., 2007), and ethical issues in the case of ESCs (Zacharias et al., 2011).

As an alternative, we and other group recently developed chemically derived hepatic progenitors (CdHs) from primary human (Kim et al., 2019b) or mouse (Katsuda et al., 2017) hepatocytes. CdHs have the capacity for rapid proliferation after reprogramming, can be stably cultured over 10 passages, exhibit adequate differentiation into hepatocytes and biliary epithelial cells, and can repopulate the liver in model disease mice after transplantation. Here, we generated mouse CdHs (mCdHs) from HT1 mouse hepatocytes by the addition of relevant chemical compounds and used the cells for *ex vivo* gene editing therapy. We corrected the *Fah* gene mutation using an ABE or a prime editing system (Anzalone et al., 2019), which was transfected into the cells through electroporation, a non-viral method. The corrected mCdHs were transplanted into the liver of HT1 model mice, and the transplanted mice survived even after NTBC withdrawal, indicating that our *ex vivo* gene editing strategy could be a suitable and reliable tool for clinical applications in hereditary liver diseases.

## Results

To generate mCdHs from HT1 model mice, we examined whether the previous protocol used for human hepatocyte reprogramming could also be applied to PHs from mice (HT1-mPHs). To this end, we treated the PHs with one growth factor and two chemical compounds (referred to as HAC), which include hepatocyte growth factor (HGF), A83-01 (TGF-β inhibitor), and CHIR99021 (GSK-3 inhibitor) (**Figure 1A**). Interestingly, we found that HAC-treated HT1-mPHs exhibited a small epithelial cell morphology three days after treatment and that the cell populations expanded to cover the dishes after eight days (**Figure 1B**). We confirmed that those proliferating cells expressed hepatic stem-/progenitor-specific markers including Krt19, Sox9, and Afp (**Figures 1B, S1A** and **S1B**); thus, we defined the cells as chemically derived hepatic progenitors from HT1 mice (HT1-mCdHs). The HT1-mCdHs showed no significant differences in gene expression levels or proliferation capacity compared with chemically derived hepatic progenitors from a wild-type C57BL/6N mouse (WT-mCdHs) (**Figure S1A and S1C**). The HT1-mCdHs were passaged stably through passage 21, indicating that they function as a stable cell line that could produce gene corrected clones (**Figure S1D**).

**Figure 1.**
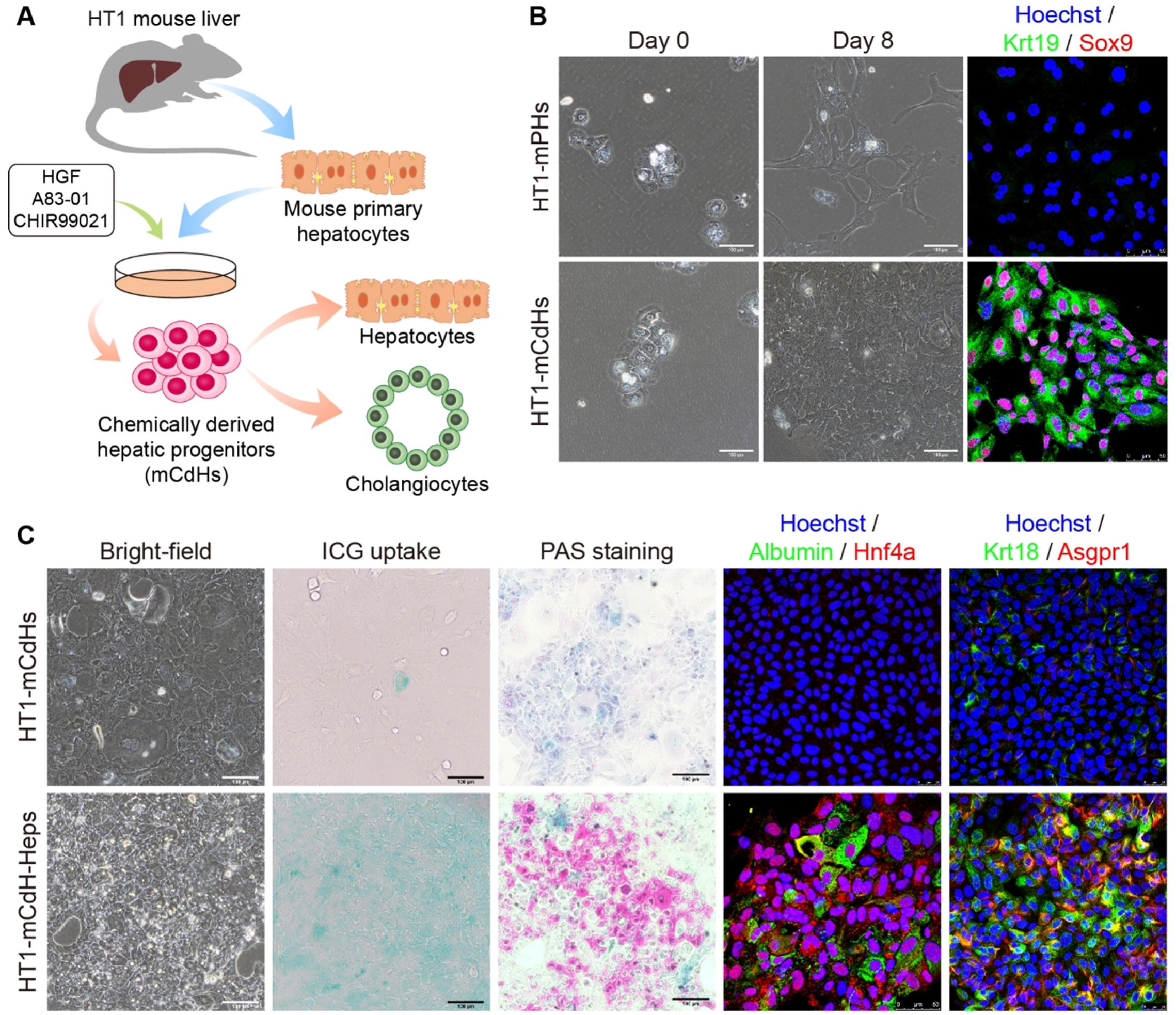
Generation and characterization of HT1-mCdHs. (A) Schematic diagram of the reprogramming method used to generate chemically derived hepatic progenitors from the HT1 mouse. (B) Freshly isolated HT1 mouse primary hepatocytes (PH) were cultured in reprogramming medium for eight days in the absence or presence of HAC. Immunofluorescence staining of hepatic progenitor markers Krt19 and Sox9 is shown. Nuclei were counterstained with Hoechst 33342. Scale bars, 100 μm and 50 μm. (C) Characteristics of HT1-mCdHs after culture under hepatic differentiation conditions, when they exhibit a mature hepatocyte phenotype. Bright-field, ICG uptake, PAS staining, and immunofluorescence staining for mature hepatocyte-specific markers are shown. Nuclei were counterstained with Hoechst 33342. Scale bars, 100 μm and 50 μm.

Hepatic progenitor cells have the capacity to differentiate into both mature hepatic cells and cholangiocytes. To examine the differentiation capacities of HT1-mCdHs, we first cultured them under hepatic differentiation conditions. We found that the hepatocyte-like cells differentiated from HT1-mCdHs (HT1-mCdHs-Heps) had acquired both a mature hepatocyte morphology and mature hepatic characteristics, as shown by analysis of indocyanine green (ICG) uptake and periodic acid-Schiff (PAS) staining (**Figure 1C**). Immunofluorescence staining showed that mature hepatocyte-specific markers, including albumin, Hnf4a, Krt18, and Asgpr1, were expressed after hepatic differentiation (**Figure 1C**), indicating that the mCdHs can re-differentiate into mature hepatocytes under the proper conditions. These characteristics were confirmed by quantitative real-time PCR (qRT-PCR) (**Figure S2A**). In addition, we conducted another experiment, involving three-dimensional culture methods, in which we induced HT1-mCdHs to differentiate into cholangiocytes. The resulting differentiated cells (HT1-mCdHs-Chols) formed characteristic tubular-like structures (**Figure S2B**) and expressed higher levels of the cholangiocytic-specific markers *Krt19, Cftr, Ae2*, and *Aqpr1* than did HT1-mCdHs and HT1-mPHs (**Figure S2C**). Taken together, these results show that we successfully established HT1-mCdHs that have a bipotent capacity to differentiate to both hepatocytes and cholangiocytes.

We next sought to establish a precise gene correction strategy for the HT1-mCdHs. The HT1 model mouse has a G>A point mutation at the 3’ end of *Fah* exon 8 that causes exon 8 skipping during the splicing process, resulting in production of non-functional FAH enzyme (**Figure 2A**). As a means of correcting the pathogenic mutation in HT1-mCdHs, we tested both ABEs and prime editors (PEs). We first designed a single-guide RNA (sgRNA) for use with a previously developed version of ABE, ABEmax, which recognizes an NGG protospacer adjacent motif (PAM) (Koblan et al., 2018). In this approach, the point mutation was positioned near but not in the editing window (4^th^-7^th^). We transfected the ABEmax-encoding plasmid together with the sgRNA-encoding plasmid into the HT1-mCdHs via electroporation; three days later, base editing outcomes in bulk cell populations were assessed by high-throughput sequencing. The results showed that the adenosine at the position at which the change was desired (A9) underwent base conversion with an efficiency of 1.6%, whereas a bystander A (A6) was more efficiently converted, with 29.4% efficiency (**Figures 2B** and **S3A**), which is an expected result because ABEmax more readily edits the sixth versus the ninth position. With the aim of reducing the frequency of this bystander base conversion, we tested a version of ABEmax (NG-ABEmax) that recognizes an NG PAM (Jeong et al., 2019), but found that the desired adenosine (now A3) was rarely converted (0.2%) (**Figures 2B** and **S3B**). To increase the editing efficiency, we then modified a recently developed ABE variant (ABE8e), which has faster deamination kinetics, so that it recognized an NG PAM (NG-ABE8e) (Richter et al., 2020). When the HT1-mCdHs were treated with NG-ABE8e, the conversion rates of the adenosine at the desired position (A3) were substantially improved, to 11%, and bystander As (A6 and A7) in intron sites were also converted at high rates (15% and 14%) (**Figures 2B** and **S3C**). Finally, we tested PEs to see if we could achieve highly precise base conversion without any bystander effects. As expected, analysis of the PE-treated HT1-mCdHs showed that the desired adenosine was converted (2.7%) without any bystander effects (**Figures 2B** and **S3D-S3F**). Taken together, our results show that we have established various ABE- and PE-based mutation correction strategies in HT1-mCdHs.

**Figure 2.**
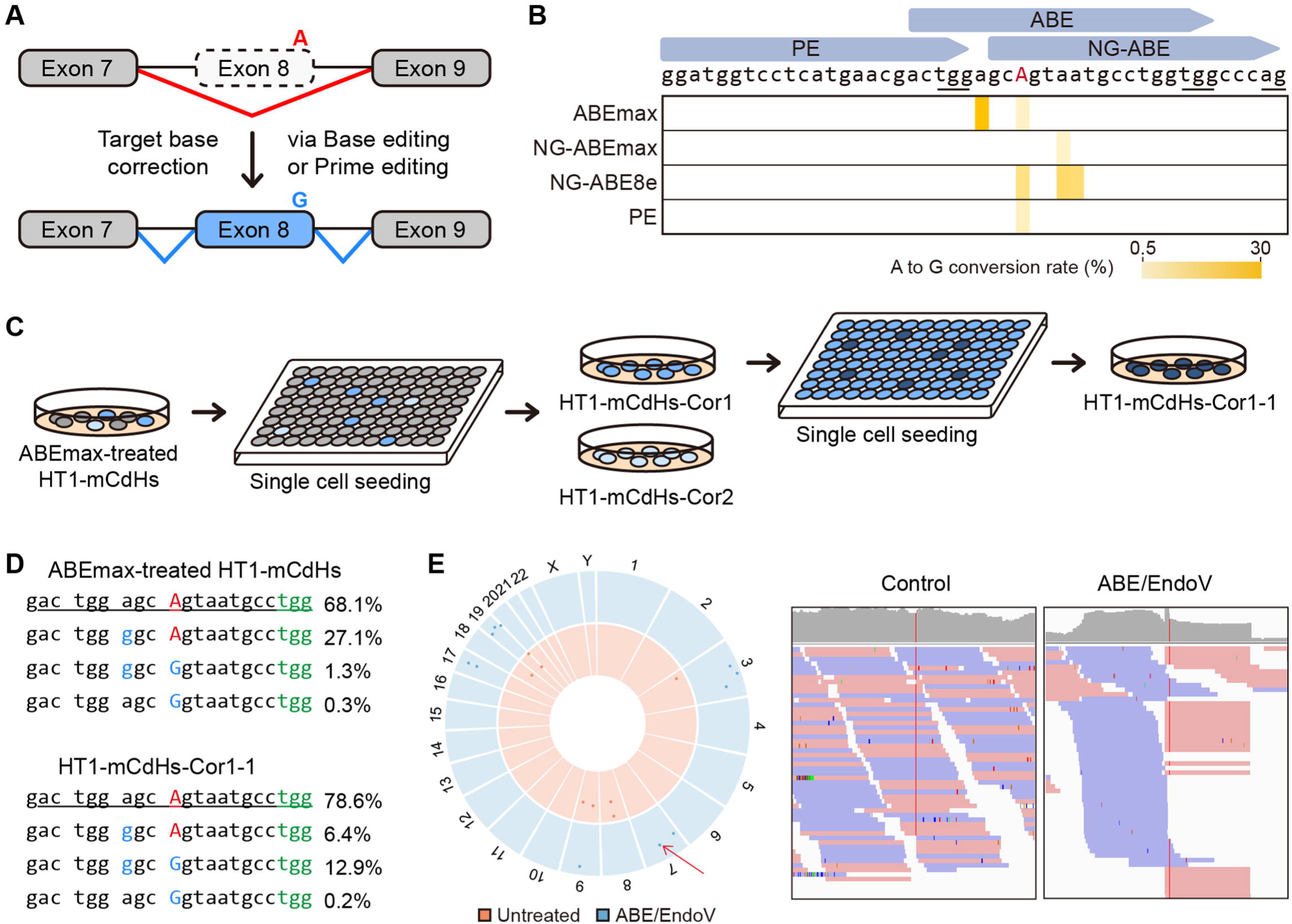
Gene editing of HT1-mCdHs with various ABE and PE systems. (A) Schematic of the base editing procedure used to correct the pathogenic mutation in HT1-mCdHs. (B) Heatmap visualizing A to G conversion rates. Blue arrows indicate the target sites, and PAM sequences are underlined. (C) Isolation of clonal cell lines containing the corrected *Fah* gene. ABE-treated HT1-mCdHs were seeded into a 96-well plate (1 cell per well), and two cell lines were selected (HT1-mCdHs-Cor1 and -Cor2). HT1-mCdHs-Cor1 cells were again seeded into a 96-well plate, to obtain HT1-mCdHs-Cor1-1. The base editing efficiency in each cell line was confirmed by high-throughput sequencing. (D) Sequences at the target site and the proportion of each sequence in ABEmax-treated HT1-mCdHs (top) and HT1-mCdHs-Cor1-1 (bottom) analyzed by high-throughput sequencing. The wild-type (WT) sequence is underlined, the pathogenic mutation is colored in red, the edited sequences in blue, and the PAM sequence in green. (E) Genome-wide Circos plot showing DNA cleavage scores from Digenome-seq analysis (left). Cleavage scores obtained from intact genomic DNA and digested DNA treated with ABE and Endo V are shown in orange and blue, respectively. The on-target site is noted by a red arrow. A representative Integrative Genomics Viewer image at the on-target site (right). After the ABE and Endo V treatment, many sequence reads near the on-target site showed a straight pattern.

For further analysis, we used ABEmax-treated HT1-mCdHs. To isolate clonal cell lines that contain the corrected *Fah* gene, we diluted the bulk population of ABE-treated cells and obtained two corrected clonal cell lines, named HT1-mCdHs-Cor1 and HT1-mCdHs-Cor2 (**Figure 2C** and **Table S1**). Notably, each cell line was associated with at least four different sequence patterns at the site of interest. To exclude the possibility that these lines were not clonal, we diluted the HT1-mCdHs-Cor1 cell population again to isolate single cells and observed that all resulting clones had at least four different sequence patterns, suggesting that the HT1-mCdHs might be polyploid, similar to hepatocytes. Flow cytometry analysis confirmed the polyploid nature of mCdHs (**Figure S3G**). From the second set of clones, we selected the cell line (named HT1-mCdHs-Cor1-1) with the highest frequency of the desired corrected sequence (13.1%) (**Figure 2D**). To examine ABEmax-mediated genome-wide off-target effects in HT1-mCdHs-Cor1-1, we performed restriction EndoV enzyme coupled Digenome-seq (Kim et al., 2019a), a method for identifying *in vitro* off-target cleavage sites, which indicated that such cells have eleven *in vitro* cleavage sites, including the on-target site (**Figure 2E** and **Table S2**). However, none of the off-target sites identified *in vitro* were reliably cleaved *in vivo*.

We next tested whether the corrected mCdHs would exhibit a reliable repopulation capacity, and thus therapeutic potential, in HT1 mice. We performed intrasplenical transplantation of the partially corrected HT1-mCdHs-Cor1-1 line, which was confirmed to lack significant off-target effects, into HT1 mice. Seven days before transplantation, NTBC was withdrawn from the drinking water of nine HT1 mice so that liver damage would be induced, thereby facilitating transplantation of the HT1-mCdHs-Cor1-1 cells (**Figure 3A**). Phosphate buffered saline (PBS) and HT1-mCdHs cells were used as negative controls, and PHs from wild-type mice (WT-mPHs cells) were used as a positive control. After transplantation, mice from the the PBS injected (5 mice) and HT1-mCdHs treated (5 mice) groups rapidly died; all mice were dead by day 90 (**Figure 3B**). Furthermore, all animals (5 mice) in the WT-mPHs treated group died at around 120 days. However, to our surprise, two from the HT1-mCdHs-Cor1-1 treated group (9 mice) survived for more than 180 days, indicating a fundamental treatment-induced improvement of HT1 disease in the absence of NTBC. In the case of the mice that survived for more than 180 days, levels of serum biomarkers including aspartate transaminase (AST), alanine transaminase (ALT), total bilirubin, and albumin (ALB) showed that liver damage was significantly decreased after transplantation of HT1-mCdHs-Cor1-1 cells (**Figure 3C**).

**Figure 3.**
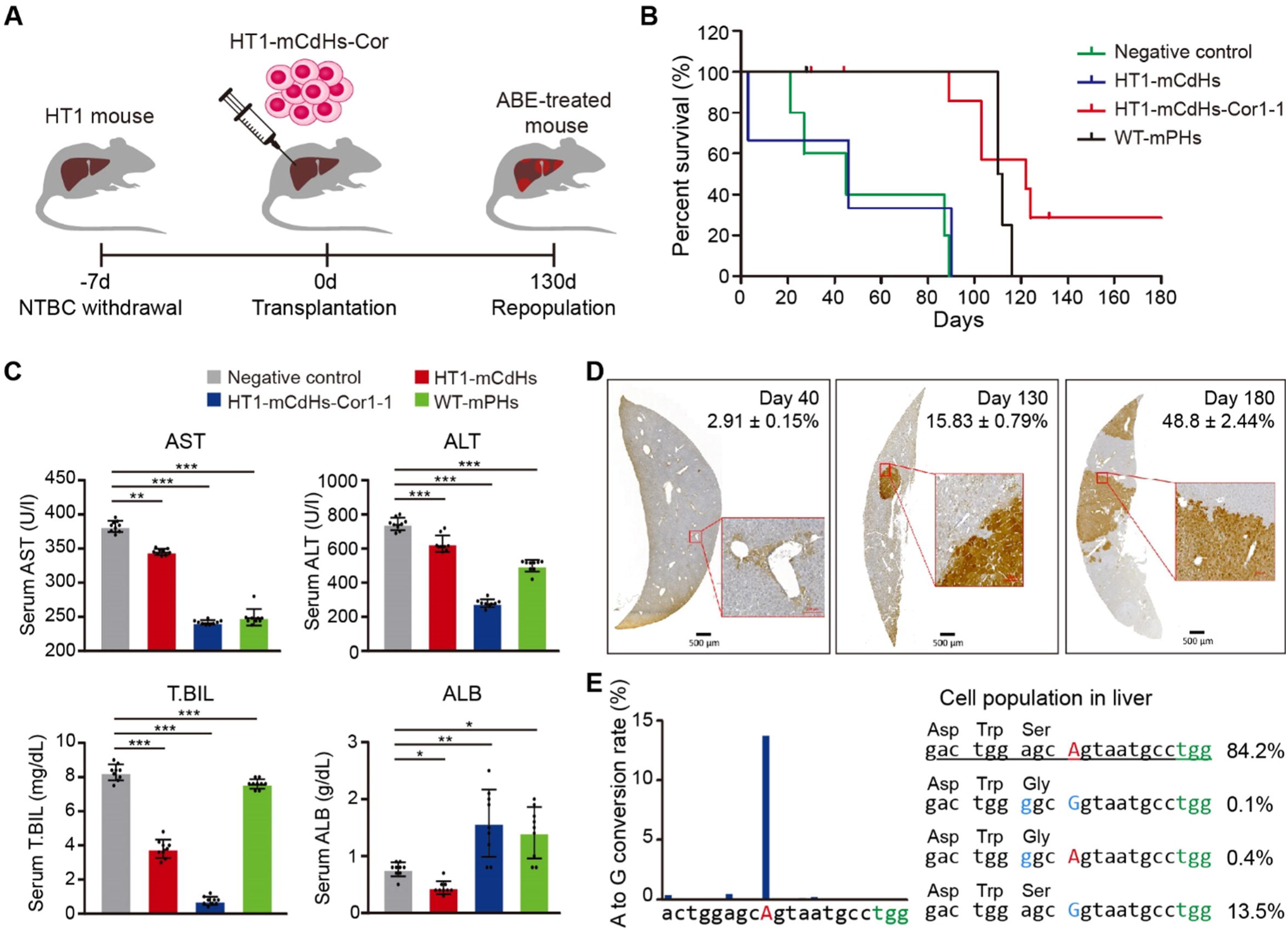
Therapeutic effects of HT1-mCdHs-Cor cells in a mouse model of HT1.(A) Scheme of HT1-mCdHs-Cor transplantation into the HT1 mouse model. (B) Survival curves of HT1 mice with or without cell transplantation. Negative control (PBS injected, 5 mice); HT1-mCdHs (5 mice); HT1-mCdHs-Cor1-1 (9 mice); and WT-mPHs (5 mice). (C) Serum levels of AST, ALT, total bilirubin, and ALB in the negative control, HT1-mCdHs, HT1-mCdHs-Cor1-1, and WT-mPHs groups. Data were analyzed by *t* test, \**p*<0.05, \*\**p*<0.01, \*\*\**p*<0.001. (D) Immunohistochemical staining of FAH in the liver at day 40, 130, and 180 after transplantation of HT1-mCdHs-Cor1-1. Scale bars, 500 μm. (E) A to G conversion rate (left) and proportion of each sequence at the target site (right) in liver tissue from HT1 mice at day 180 after transplantation with HT1-mCdHs-Cor1-1. Color code is as in Figure 2D.

To confirm the repopulation capacity of HT1-mCdHs-Cor1-1 cells, we examined FAH-positive cell populations in the mice from the HT1-mCdHs-Cor1-1 transplanted group at 40, 130 and 180 days. FAH-positive cell populations were found engrafted around the liver veins at 40 days after transplantation (**Figure 3D**). After 130 days, the area occupied by FAH-positive cells had increased to 15% of the liver slice, and further increased to almost 50% at 180 days, and these cells showed a different morphology than the original PHs (**Figures 3D**). Furthermore, it is notable that the frequency of alleles in which the target adenosine (A9) had been precisely edited without the bystander editing had abruptly increased (from 0.2 to 13.3%) by day 180, whereas the frequency of alleles in which both of the target (A9) and the bystander adenosines (A6) had been edited was decreased (from 12.9 to 0.1%) in the HT1-mCdHs-Cor1-1 treated mice (**Figure 3E**), indicating that cells containing the corrected alleles became dominant in the liver during cell duplication, similar to observations in previous studies (Song et al., 2020). Conversion to A6 to G will lead an amino acid mutation (serine to glycine, S235G) near the FAH enzyme active site (D233, K234), which would impede FAH enzyme function; thus, cells possessing an A6 substitution might be eliminated during NTBC withdrawal *in vivo*.

To investigate the reproducibility of our *ex vivo* gene editing strategy, we repeated the experiments with other corrected mCdHs lines, HT1-mCdHs-Cor1 and HT1-mCdHs-Cor2 (**Figure S4A**). We confirmed that the mice in the HT1-mCdHs-Cor1 (4 mice) and HT1-mCdHs-Cor2 (7 mice) treated groups also survived for more than 130 days under NTBC withdrawal conditions (**Figure S4B**). In addition, levels of markers that indicate liver damage were decreased (**Figure S4C**). Likewise, FAH-positive cell populations in these two groups were observed to show similar patterns as those in the HT1-mCdHs-Cor1-1 transplated group (**Figures S4D and S4E**), although the frequency of sequences in which the mutation had been corrected was lower than in the HT1-mCdHs-Cor1-1 group. These results indicate that our *ex vivo* gene editing strategy is a reliable and solid approach for HT1 disease treatment in mice.

## Discussion

Human genetic disorders are often associated with severe pathological phenotypes, but few curative therapies are available. In this study, we successfully generated mCdHs from HT1 mice and corrected the pathogenic *Fah* gene mutation in these cells using both base editing and prime editing tools that were delivered via a non-viral, electroporation method. Further, we demonstrated that the corrected mCdH population would expand in the liver after transplantation, allowing the mice to survive even under NTBC withdrawal conditions. Our *ex vivo* gene editing strategy, involving base correction in mCdHs, has several important advantages compared to previous strategies: i) exogenous genetic factors, which can cause unexpected genetic changes, are not required for the induction of hepatic progenitors from PHs. ii) we can assess the mutation correction rates and genome-wide off-target effects in the cells with a high degree of accuracy prior to transplantation, so we can select a cell line in which base correction was more efficient and that exhibits negligible off-target effects. iii) the gene editing efficiency is less critical than that in *in vivo* gene correction strategies because corrected cells can be enriched *in vitro*, and non-viral delivery of CRISPR tools is easily accomplished.

In this study, we established various gene correction strategies in mCdHs that involved several gene editing systems, including the canonical ABE that recognizes an NGG PAM (i.e., ABEmax), an ABE that recognizes an NG PAM (i.e., NG-ABE8e), and prime editing. Although we demonstrated *ex vivo* therapeutic treatment only with ABEmax-treated mCdHs, showing that mice transplanted with such corrected cells survived under NTBC withdrawal conditions, NG-ABE8e-treated and PE-treated mCdHs should also improve symptoms in a similar way. These two editing systems have advantages compared to the ABEmax. In particular, NG-ABE8e induced bystander mutations in intron sites and PE did not induce detectable bystander mutations. Therefore, highly precise gene correction will be made possible by using those tools in the future. Our novel *ex vivo* therapeutic editing strategy opens a new avenue for treating not only HT1 disease, but other genetic diseases that affect the liver. Small molecule-mediated cellular reprogramming and gene correction techniques provide the potential for new clinical applications in various cell types.

## Acknowledgments

This research was supported by grants from the National Research Foundation of Korea (NRF) no. 2018M3A9H3022412 and Korea Healthcare Technology R&D Project (HI16C1012) to S.B. and by Medical Research Center (NRF-2017R1A5A2015395) and Basic Science Research Program (2019R1F1A106114812) to D.C.

## Author Contributions

S.B. and D.C conceived this project. Y. K., J. Y., and S.A.H. performed and analyzed most of the experiments. J.E. and K.J assisted with histological analysis. S.-N.L and J.-S.W performed protein works. Y.K. and S.B. wrote the manuscript with the approval of all other authors. J.J., S.B., and D.C. supervised the research.

## Declaration of Interests

Y.K., J.Y., S.-A.H., J.J., S.B., and D.C. have filed a patent application based on this work.

## STAR★Methods

### LEAD CONTACT AND MATERIALS AVAILABILITY

Further information and requests for resources and reagents should be directed to and will be fulfilled by the Lead Contact, Sangsu Bae.

### EXPERIMENTAL MODEL AND SUBJECT DETAILS

#### Animals

HT1 mice were a gift from Hyoungbum (Henry) Kim. Experiments were performed on 6-8-week-old male and female mice. The mice were housed and cared for under specific pathogen-free conditions in accordance with the Principles of Laboratory Animal Care and the Guide for the Use of Laboratory Animals of HYU Industry-University Cooperation Foundation regulations (2018-0196A). Liver damage was induced in HT1 mice by withdrawal of NTBC for 1 week.

#### Isolation of primary hepatocytes and cell culture

To isolate *Fah*^*-/-*^mouse primary hepatocytes, livers in HT1 mice were perfused through the portal vein with solution A (0.19 g/L EDTA (Sigma-Aldrich), 8 g/L NaCl, 0.4 g/L KCl, 0.078 g/L NaH_2_PO4·2H_2_O, 0.151 g/L Na_2_HPO4·12H_2_O, and 0.19 g/L HEPES) for 5 min at 37°C, followed by solution B (0.3 g/L collagenase (Worthington Biochemical), 0.56 g/L CaCl_2_, 8 g/L NaCl, 0.4 g/L KCl, 0.078 g/L NaH_2_PO4·2H_2_O, 0.151 g/L Na_2_HPO4·12H_2_O, and 0.19 g/L HEPES) for 8 min at 37°C. Viable primary hepatocytes were obtained by isodensity centrifugation in Percoll solution (GE Healthcare). Isolated *Fah*^*-/-*^mouse primary hepatocytes were seeded in a collagen-coated dish at 2,000 cells/cm^2^. Cells were then cultured in William’s E medium (Gibco) in a humidified atmosphere containing 5% CO_2_ at 37°C.

For generating chemically derived hepatic progenitors from the HT1 primary hepatocytes, 1 day after seeding the medium was changed to reprogramming medium [DMEM/F-12 medium containing 1% fetal bovine serum (FBS) (Gibco), 1% insulin-transferrin-selenium (Gibco), 0.1 μM dexamethasone (Sigma-Aldrich), 10 mM nicotinamide (Sigma-Aldrich), 50 μM β-mercaptoethanol (Sigma-Aldrich), 1% penicillin/streptomycin (Gibco), 20 ng/mL epidermal growth factor (Peprotech), 20 ng/mL hepatocyte growth factor (Peprotech), 4 μM A83-01 (Sigma-Aldrich), and 3 μM CHIR99021 (Sigma-Aldrich)]. The reprogramming medium was changed every couple of days. Every 4 to 6 days, the cells were passaged by first dissociating them from the plates by treatment with 1X TrypLE Express Enzyme (Gibco), diluting the released cells into fresh medium at a ratio of 1:4, and plating them in fresh collagen-coated dishes. After base editing, the bulk population of cells was diluted and seeded into 96-well plates so that single cell-derived clones could be picked.

For hepatic differentiation, HT1-mCdHs were seeded on collagen-coated dishes at 1,000 cells/cm^2^. After a 1-day incubation, the medium was changed to differentiation medium consisted of the reprogramming medium supplemented with 20 ng/mL oncostatin M (Prospec) and 10 μM dexamethasone; the medium was changed every two days thereafter. After 6 days, the cells were covered with Matrigel (Corning) diluted with differentiation medium at a 1:7 ratio and cultured 2 more days.

For cholangiocytic differentiation, HT1-mCdHs were harvested by treatment with 1X TrypLE Express Enzyme and resuspended at a density of 1 × 10^5^ cells/well in 6-well plates in DMEM/F-12 medium containing 10% FBS and 20 ng/mL hepatocyte growth factor [designated cholangiocyte differentiation medium (CDM)]. The CDM was mixed on ice with an equal volume of collagen type I (pH 7.0) and incubated for 30 min at 37°C for solidifying. Then, the cells were overlaid with mixtures and cultured for 7 days. The medium was changed every two days.

## METHOD DETAILS

### Immunostaining

For immunocytochemistry, the cells were fixed in 4% paraformaldehyde at 4°C overnight. The fixed cells were washed in PBS and then treated with PBS containing 0.2% Triton X-100 for 10 min at room temperature. Next, cells were treated with blocking solution consisting of 1% bovine serum albumin, 22.52 ng/mL glycine, and 0.1% Tween 20 in PBS for 1 hr at room temperature, after which the cells were incubated with primary antibodies diluted in blocking solution at 4°C overnight. After washing, the primary antibodies were detected using Alexa Fluor 488- or 594-conjugated secondary antibodies (Thermo Fisher Scientific). Nuclei were counterstained with Hoechst 33342 (1:10,000, Molecular Probes). Primary antibodies used in this study are listed in **Key Resources Table**. Stained cells were visualized under a TCS SP5 confocal microscope (Leica).

For immunohistochemistry, liver tissue samples were fixed in 10% formalin and embedded in paraffin. Sections were subjected to immunohistochemical staining. Immunohistochemical staining was performed using a Dako REAL™ EnVision™ Detection System (Dako). Anti-FAH antibody (Yecuris, 20-0034) was used as the primary antibody and nuclei were counterstained with hematoxylin. Stained tissues were viewed under a Virtual Microscope Axio Scan.Z1 (Zelss).

### Isolation of mRNA and RT-PCR analysis

Total RNAs were isolated using Trizol Reagent (Gibco). Then, 1 μg RNA samples were reverse transcribed using a Transcriptor First Strand cDNA Synthesis Kit (Roche). RT-PCR was performed using a CFX Connect Real-Time PCR Detection system (Bio-Rad); each reaction contained 10 μL of qPCR PreMix (Dyne Bio), 1 μL of cDNA, and oligonucleotide primers. Reactions were analyzed in triplicate for each gene. The PCR cycles consisted of 40 cycles of 95°C for 20 sec followed by 60°C for 40 sec. Melting curves and melting peak data were obtained to characterize the PCR products. The primer sequences are listed in **Table S3**.

### Calculation of the doubling time

HT1-mCdHs were seeded at a density of 1 × 10^4^ cells/well onto collagen-coated 6-well plates. Cell numbers were determined on day 3 and 7. The doubling time was calculated using the following formula:

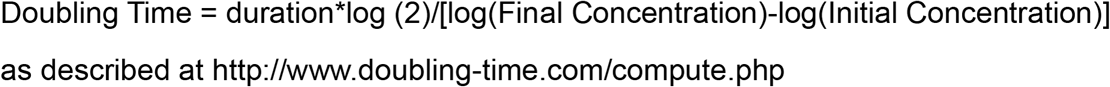

### PAS staining ICG uptake

To detect glycogen, cells were stained with PAS reagent using a PAS staining kit (Abcam) in the presence or absence of diastase (Sigma) as recommended by the supplier. To detect ICG (Sigma) uptake, cells were incubated in medium containing 1 mg/mL of ICG for 30 min at 37°C and examined under a phase-contrast microscope.

### Construction of sgRNA- and pegRNA-expressing plasmids

To construct sgRNA-expressing plasmids, complementary oligos representing the target sequences were annealed and cloned into pRG2 (Addgene #104174). To construct pegRNA-expressing plasmids, complementary oligos representing the target sequences, sgRNA scaffold, and 3’ extensions were annealed and cloned into pU6-pegRNA-GG-acceptor (Addgene #132777).

### Transfection of HT1-mCdHs

Electroporation was performed using an Amaxa 4-D device (Lonza) or a Neon Transfection System (Thermo Fisher). For the Amaxa 4-D device, a P3 Primary Cell 4D-Nucleofector X Kit (program EX-147) was used. 200,000 HT1-mCdHs were electroporated with 750 ng of ABEmax-encoding plasmid (Addgene, #112095) and 250 ng of sgRNA-encoding plasmid. Using the Neon Transfection System, 100,000 HT1-mCdHs were transfected with 900 ng of PE2-encoding plasmid (Addgene #132775), 300 ng of pegRNA-encoding plasmid, and 83 ng of nicking guide RNA-(ngRNA-) encoding plasmid or 900 ng of NG-ABE-encoding plasmid (NG-ABE8e, Addgene #138491) and 250 ng of sgRNA-encoding plasmid with the following parameters: voltage, 1,200; width, 50 ms; number, 1. NG-ABEmax-encoding plasmid wasconstructed in our lab based on the appropriate backbone plasmids (Addgene # 112095).

Three days after transfection, the cells were harvested by centrifugation in preparation for high-throughput sequencing. The cell pellet was resuspended in 100 μl of Proteinase K extraction buffer [40 mM Tris-HCl (pH 8.0) (Sigma), 1% Tween-20 (Sigma), 0.2 mM EDTA (Sigma), 10 mg of proteinase K, 0.2% nonidet P-40 (VWR Life Science)], incubated at 60°C for 15 min, and heated to 98°C for 5 min.

### High-throughput sequencing

ABE target sites were amplified from extracted genomic DNA using SUN-PCR blend (Sun Genetics). The PCR products were purified using Expin™ PCR SV mini (GeneAll) and sequenced using a MiniSeq Sequencing System (Illumina). The results were analyzed using Cas-Analyzer (http://www.rgenome.net/cas-analyzer/) and BE-Analyzer (http://www.rgenome.net/be-analyzer/). The primers are listed in **Table S3**.

### Digenome-seq

Genomic DNA was extracted from HT1-mCdHs using a DNeasy Blood & Tissue Kit (Qiagen). 8 μg of the genomic DNA was incubated with 32 μg of ABE pre-incubated with 24 μg of *in vitro* transcribed sgRNA at room temperature for 5 min, after which 300 μL of 2X BF buffer (Biosesang) was added and the reaction volume brought to 600 μL. That mixture was incubated at 37°C for 16 h. After RNase A (50 μg/mL, Thermo Scientific) treatment at 37°C for 15 min, the ABE-treated genomic DNA was purified using a DNeasy Blood & Tissue Kit (Qiagen). 3 μg of the purified DNA was digested with 8 units of Endo V (New England Biolabs) in a 200 μL reaction at 37 °C for 2 h. The genomic DNA was then purified using a DNeasy Blood & Tissue Kit (Qiagen). Whole genome sequencing was performed with 1 μg of the digested DNA using a HiSeq X Ten Sequencer (Illumina) at Macrogen (South Korea).

### *In vitro* transcription of sgRNAs

To generate a template for *in vitro* transcription, a forward oligo containing a T7 RNA polymerase promoter and the target sequence and a reverse oligo containing a guide RNA scaffold were purchased from Macrogen (South Korea) and extended using Phusion DNA polymerase (Thermo Scientific). The extended DNA products were purified using Expin PCR SV mini (GeneAll) and transcribed by T7 RNA polymerase (New England Biolabs). After incubation at 37 °C for 16 h, DNA templates were degraded with DNase I (New England Biolabs), and the RNA products were purified with Expin PCR SV mini (GeneAll). The oligos are listed in **Table S4**.

### Transplantation

Seven days before cell transplantation into mice, NTBC was withdrawn from the drinking water. 1 × 10^6^ cells in 100 μL PBS were transplanted into the inferior pole of the spleen. NTBC was transiently given every 3 days when mice reached 80% of their original weight.

After transplantation, serum was collected for biomarker analysis. Serum was diluted at a ratio of 1:4 to obtain means.

### Analysis of ploidy

HT1 mPHs, mCdHs, and mCdHs-Cor1-1 cells were dissociated from plates by trypsinization and then incubated with 15 μg/mL of Hoechst 33342 and 5 μM of reserpine for 30 min at 37°C. Cell ploidy was analyzed with a FACSCanto II (BD Biosciences) as described by Duncan *et al*. (Duncan et al., 2010).

## QUANTIFICATION AND STATISTICAL ANALYSIS

### Statistical Analysis

Doubling time experiments and qRT-PCR were performed in triple biological replicates. Quantitative data are presented as means ± standard deviations (SDs) with inferential statistics (p-values). Survival was analyzed as a Kaplan-Meier curve using GraphPad Prism 7 (GraphPad). Statistical significance was evaluated by two-tailed t-tests with significances set at * p< 0.05, **p < 0.01, and ***p < 0.001.

## DATA AND CODE AVAILABILITY

high-throughput sequencing data have been deposited in the NCBI Sequence Read Archive database (SRA; https://www.ncbi.nlm.nih.gov/sra) under accession number SUB7884878.

## Notes

### Competing Interest Statement

The authors have declared no competing interest.

