## Supplementary for "*Ex vivo* therapeutic base and prime editing using chemically derived hepatic progenitors in a mouse model of tyrosinemia type 1"

**Table of Contents**

**Figure S1**. Hepatic progenitor features of HT1-mCdHs

**Figure S2**. Bi-potent differentiation capacity of HT1-mCdHs.

**Figure S3**. Establishment of gene correction systems.

**Figure S4**. Reproducibility study of the therapeutic potential of HT1-mCdHs-Cor1 and 2 in HT1 model mice.

**Table S1**. Isolation of clonal cell lines of Fah-corrected mCdHs.

**Table S2**. Genomic sites captured by Digenome-seq

**Table S3**. Primers for qRT-PCR (A) and high-throughput sequencing (B), related to STAR Methods.

**Table S4**. Oligos for sgRNA plasmid cloning (A), pegRNA plasmid cloning (B), ngRNA plasmid cloning (C), and sgRNA in vitro transcription (D), related to STAR Methods.

**Figure S1. Hepatic progenitor features of HT1-mCdHs.** (A) Gene expression of hepatic progenitor markers determined by RT-qPCR. Gapdh was used as an internal control. Data are mean ± SD (n=9). Data were analyzed by *t* test, **p*<0.05, ***p*<0.01, ****p*<0.001. (B) Immunofluorescence staining of hepatic progenitor markers. Nuclei were counterstained with Hoechst 33342. Scale bars, 50 μm. (C) Doubling times of WT-mCdHs and HT1-mCdHs cultured for 72 hours. Data are mean ± SD (n=3). (D) Bright-field images of HT1-mCdHs at early (p1) and long-term (p21) passages.

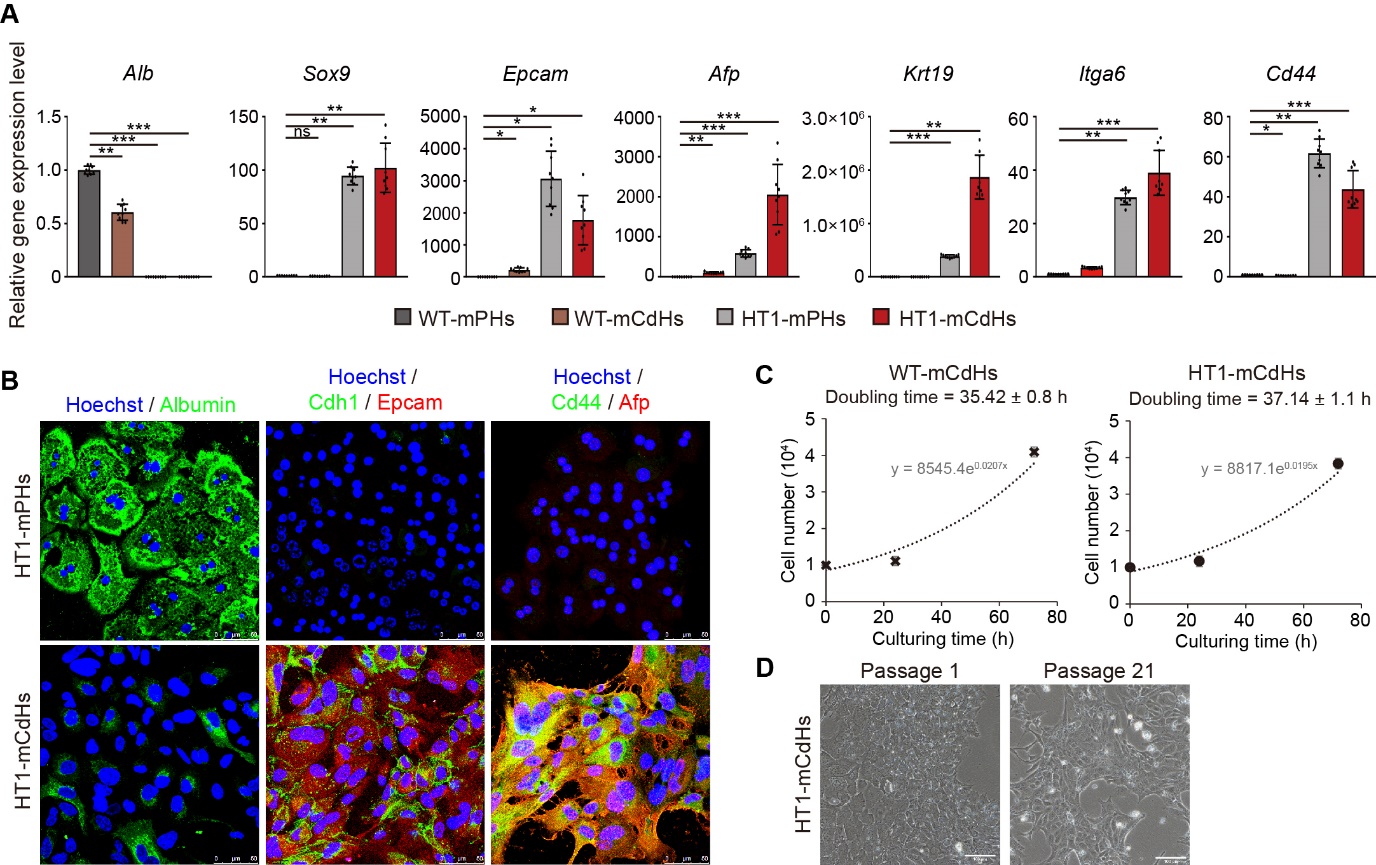

**Figure S2. Bi-potent differentiation capacity of HT1-mCdHs.** (A) Gene expression of mature hepatocyte-specific markers determined by RT-qPCR. Gapdh was used as an internal control. Data are mean ± SD (n=9). Data were analyzed by *t* test, **p*<0.05, ***p*<0.01, ****p*<0.001. (B) Bright-field image of HT1-mCdH-Chols, which have been cultured under conditions to induce cholangiocytic differentiation. Scale bars, 100 μm. (C) Cholangiocyte-specific marker expression by HT1-mCdH-Chols measured by RT-qPCR. Gapdh was used as an internal control. Data are mean ± SD (n=9). Data were analyzed by *t* test, **p*<0.05, ***p*<0.01, ****p*<0.001.

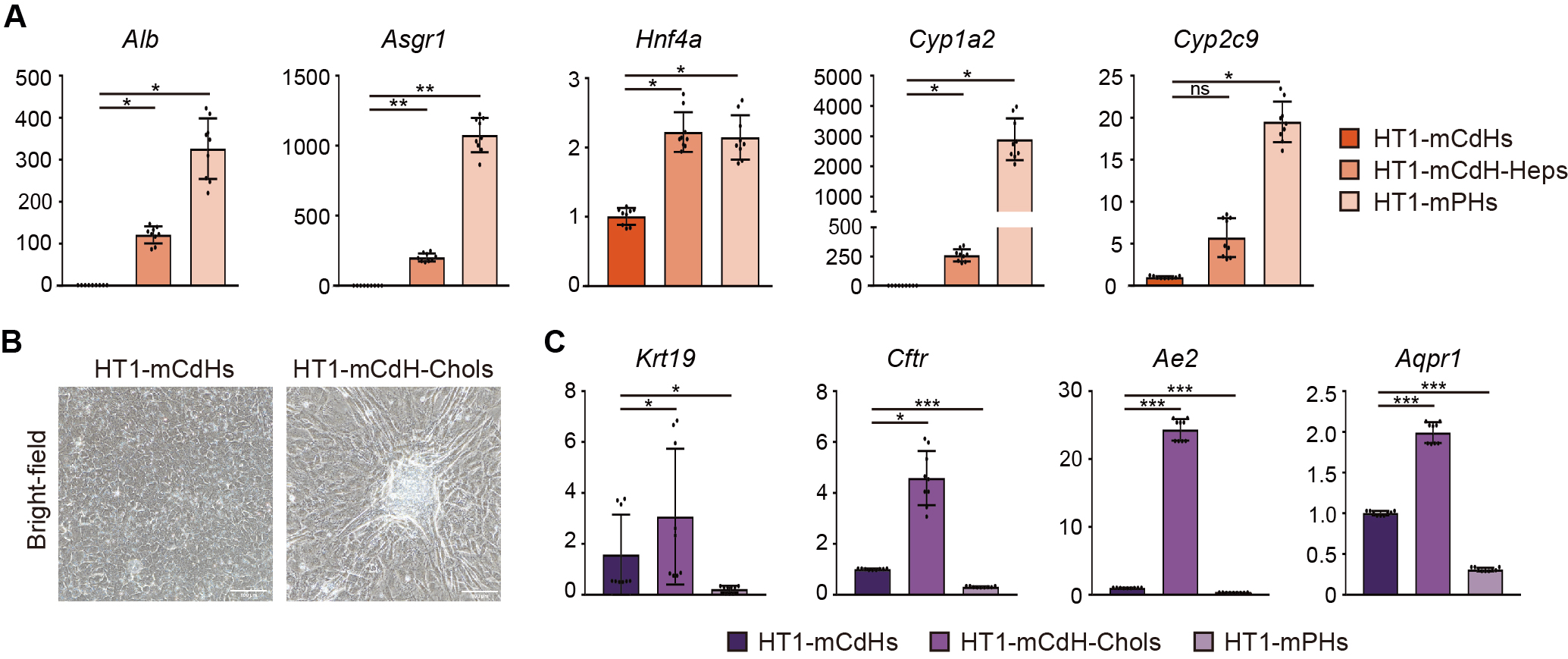

**Figure S3. Establishment of gene correction systems.** (A-C) Sequence tables showing the proportion of nucleotides at each position in the target sequence in HT1-mCdHs transfected with ABEmax (A), NG-ABEmax (B), or NG-ABE8e (C). PAM sequences are shown in green, and the G to A mutation in HT1-mCdHs is shown in red. (D) Schematic of prime editing system. In the case of prime editing target 1 (top), prime editor 3 (PE3), together with pegRNA_1 and ngRNA_1, or PE3b, together with pegRNA_1 and ngRNA_1b, were used. In the case of prime editing target 2 (bottom), PE3, together with pegRNA_2 and ngRNA_2, or PE3b, together with pegRNA_2 and ngRNA_2b, were used. pegRNA: prime editing guide RNA, ngRNA: nicking single-guide RNA. (E) The pegRNA sequence used for prime editing target 1. Primer binding site: PBS, reverse transcription template: RTT. (F) Optimization of prime editing system for target 1 by testing different PBS lengths and PEs in combination with a 15-nucleotide RTT. PE3b and PE3 showed similar editing activities but PE3 generated a higher indel (insertion and deletion) frequency (0.6%) than did PE3b. (G) Optimization of prime editing system for target 2 by testing different PBS lengths and PEs. Overall, prime editing target 2 showed lower activity than prime editing target 1. (H) Different ploidy populations of HT1 mPHs, mCdHs, and mCdHs-Cor1-1 separated by fluorescence-activated cell sorting using Hoechst 33342 fluorescence.

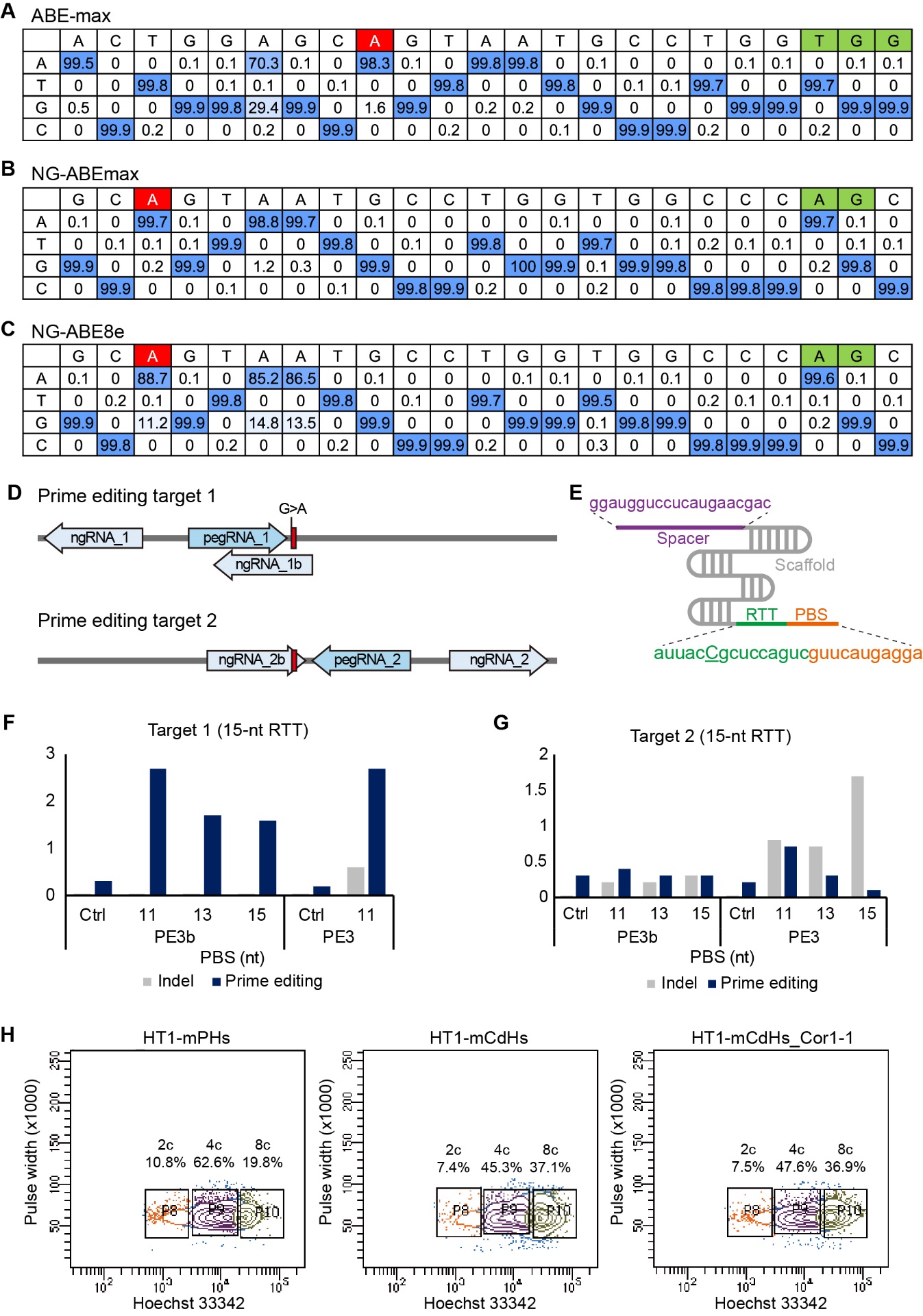

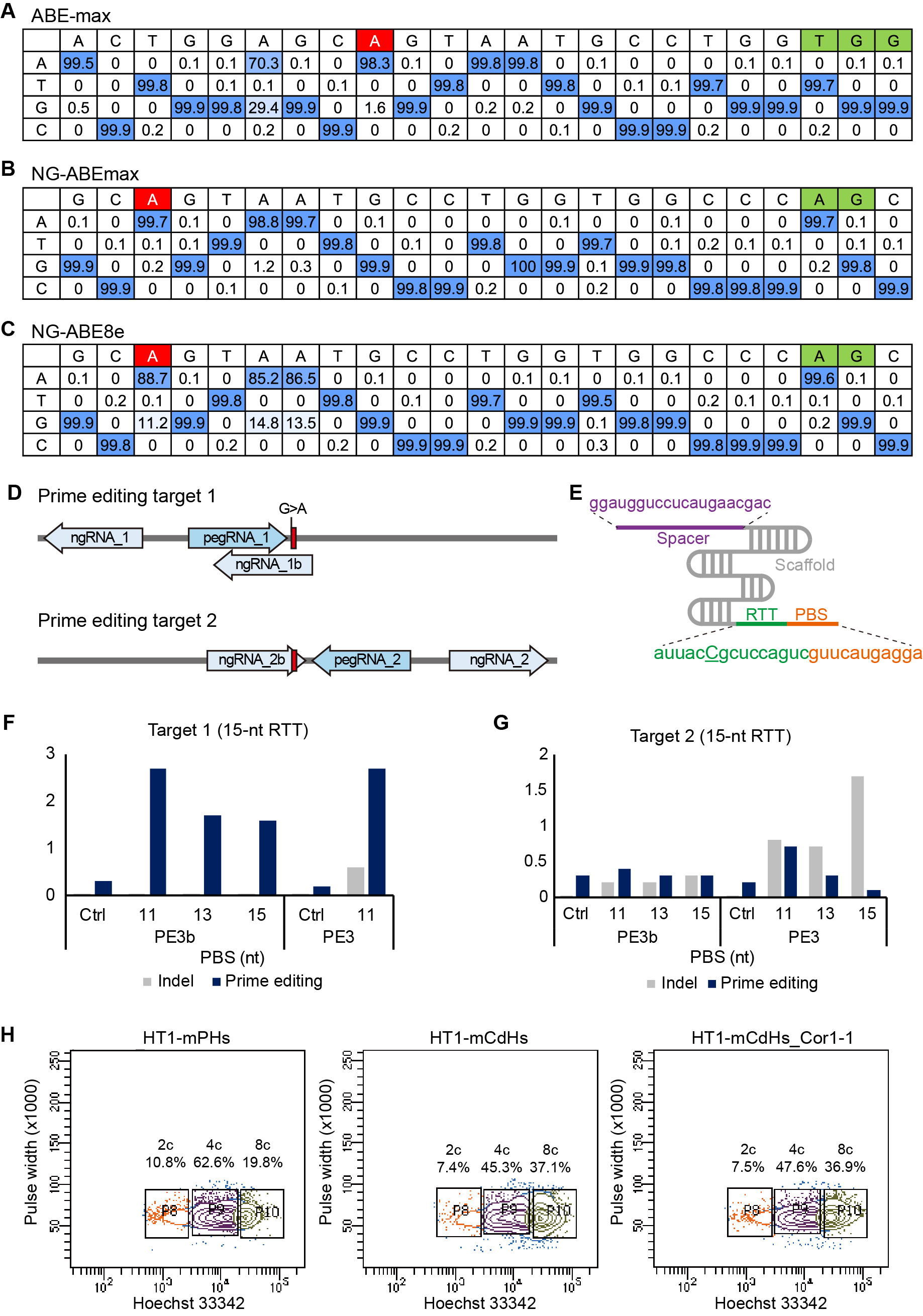

**Figure S4. Reproducibility study of the therapeutic potential of HT1-mCdHs-Cor1 and 2 in HT1 model mice.** (A) Sequences at the target site and proportion of each sequence of HT1-mCdHs-Cor1 (top) and HT1-mCdHs-Cor2 (bottom) analyzed by high-throughput sequencing. The WT sequence is underlined, the pathogenic mutation is colored in red, edited sequences in blue, and the PAM sequence in green. (B) Survival curves of HT1 mice with or without cell transplantation. Negative control (PBS injected, 5 mice); HT1-mCdHs (5 mice); HT1-mCdHs-Cor1 (4 mice); HT1-mCdHs-Cor2 (7 mice); and WT-mPHs (5 mice). (C) Serum levels of AST, ALT, total bilirubin, and ALB in the negative control, HT1-mCdHs, HT1-mCdHs-Cor1, HT1-mCdHs-Cor2, and WT-mPHs groups. Data were analyzed by *t* test, **p*<0.05, ***p*<0.01, ****p*<0.001. (D) and (E) Immunohistochemical staining of FAH and H&E staining in liver tissue from HT1 mice transplanted with HT1-mCdHs-Cor1-1 (day 180) and HT1-mCdHs-Cor 1 and 2 (day 130). Scale bars, 100 μm.

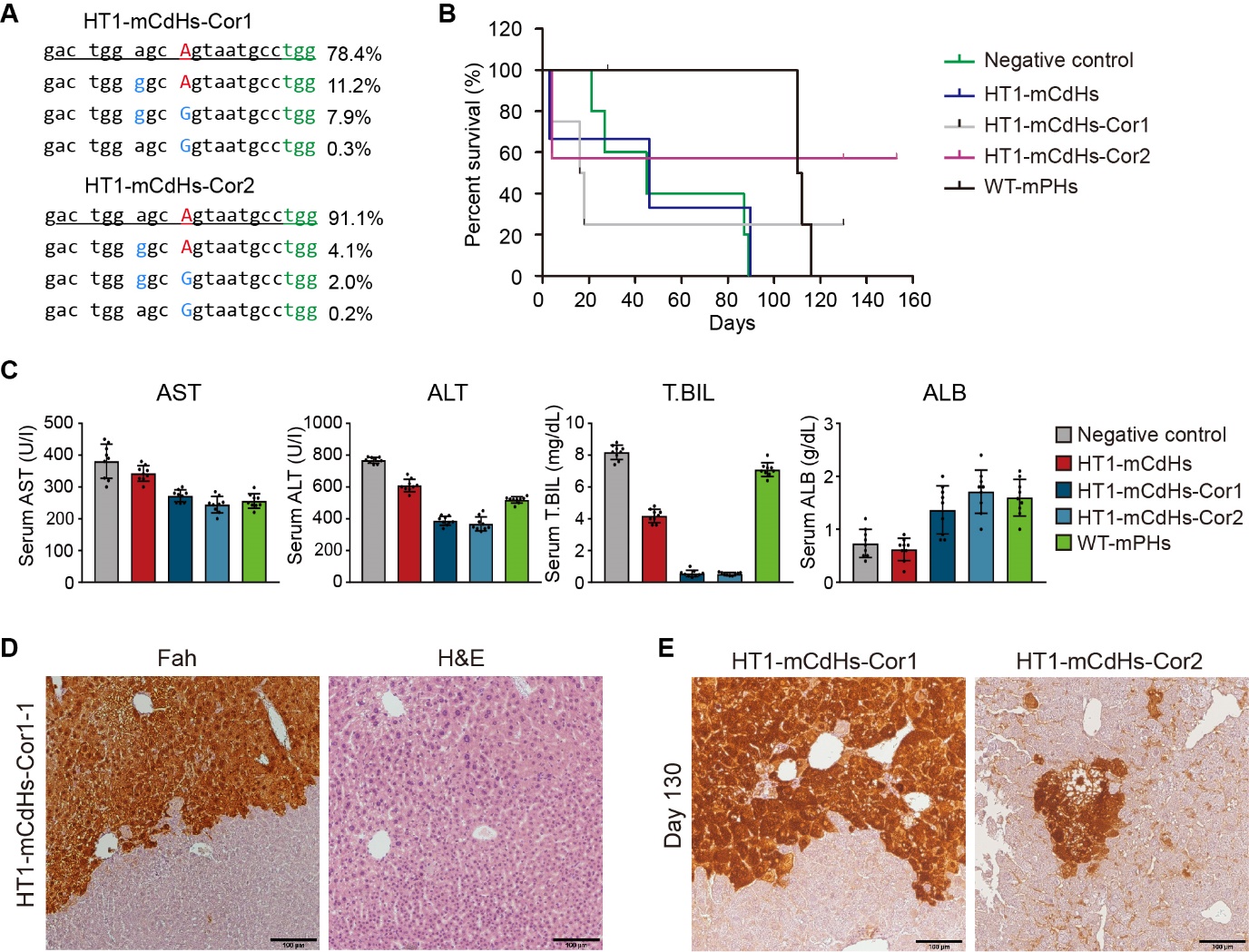

**Table S1. Isolation of clonal cell lines of *Fah*-corrected mCdHs.** Clonal cell lines were established from ABE-treated HT1-mCdHs bulk cells (A) and HT1-mCdHs-Cor1 (B).

| **Clonal cell line No.** | **Total count** | **Substitution** | | | **Indel** | |
| --- | --- | --- | --- | --- | --- | --- |
|  |  | **Count** | **Frequency at A6** | **Frequency at A9** | **Count** | **Frequency** |
| **A. Isolation of clonal cell lines from ABE-transfected HT1-mCdHs bulk cells** | | | | | | |
| S1 | 3879 | 243 | 4.0% | 0.2% | 2 | 0.1% |
| S2 | 5864 | 279 | 2.8% | 0.1% | 4 | 0.1% |
| S3 | 5513 | 2034 | 35.6% | 0.2% | 2 | 0.0% |
| S4 | 4331 | 1057 | 22.7% | 0.2% | 0 | 0.0% |
| S5 | 5762 | 1196 | 19.1% | 0.3% | 3 | 0.1% |
| S6 | 5171 | 525 | 8.5% | 0.1% | 4 | 0.1% |
| S7 | 5705 | 284 | 3.3% | 0.1% | 5 | 0.1% |
| S8 | 4718 | 1558 | 31.4% | 0.1% | 2 | 0.0% |
| S9 | 5423 | 2591 | 46.4% | 0.3% | 7 | 0.1% |
| S10 | 4342 | 304 | 5.1% | 0.3% | 6 | 0.1% |
| S11 | 5319 | 1573 | 27.7% | 0.4% | 2 | 0.0% |
| S12 | 5717 | 268 | 2.9% | 0.2% | 4 | 0.1% |
| S13 | 3818 | 171 | 2.1% | 0.2% | 2 | 0.1% |
| S14 | 5401 | 183 | 1.7% | 0.1% | 2 | 0.0% |
| S15 | 4972 | 210 | 2.3% | 0.4% | 5 | 0.1% |
| S16 | 3298 | 971 | 27.7% | 0.4% | 5 | 0.2% |
| S17 | 5279 | 747 | 12.7% | 0.3% | 4 | 0.1% |
| S18 | 5125 | 268 | 3.4% | 0.2% | 8 | 0.2% |
| S19 | 5370 | 151 | 0.9% | 0.2% | 6 | 0.1% |
| S20 | 4789 | 214 | 2.5% | 0.2% | 0 | 0.0% |
| S21  (HT1-mCdHs-Cor1) | 4794 | 1013 | 19.4% | 8.3% | 5 | 0.1% |
| S22  (HT1-mCdHs-Cor2) | 4750 | 389 | 6.1% | 2.2% | 7 | 0.1% |
| S23 | 4640 | 247 | 3.0% | 0.3% | 2 | 0.0% |
| S24 | 4561 | 188 | 2.5% | 0.2% | 5 | 0.1% |
| S25 | 3556 | 484 | 11.4% | 0.4% | 6 | 0.2% |
| S26 | 4797 | 492 | 8.4% | 0.3% | 11 | 0.2% |
| S27 | 4900 | 526 | 9.0% | 0.1% | 2 | 0.0% |
| S28 | 3354 | 129 | 2.0% | 0.1% | 3 | 0.1% |
| S29 | 4788 | 468 | 7.9% | 0.3% | 4 | 0.1% |
| S30 | 4994 | 467 | 7.7% | 0.2% | 5 | 0.1% |
| S31 | 4847 | 400 | 6.2% | 0.2% | 10 | 0.2% |
| S32 | 4588 | 163 | 1.2% | 0.2% | 0 | 0.0% |
| S33 | 5662 | 1544 | 25.6% | 0.3% | 7 | 0.1% |
| S34 | 4181 | 579 | 12.1% | 0.2% | 0 | 0.0% |
| S35 | 4925 | 1029 | 18.7% | 0.2% | 2 | 0.0% |
| S36 | 4679 | 644 | 11.9% | 1.5% | 0 | 0.0% |
| **B. Isolation of clonal cell lines from HT1-mCdHs-Cor1** | | | | | | |
| S21-1 | 3528 | 89 | 0.2% | 0.2% | 3 | 0.1% |
| S21-2 | 2576 | 65 | 0.2% | 0.2% | 0 | 0.0% |
| S21-3 | 3344 | 362 | 8.9% | 5.9% | 5 | 0.1% |
| S21-4 | 2032 | 33 | 0.1% | 0.1% | 5 | 0.2% |
| S21-5 | 3210 | 82 | 0.2% | 0.2% | 3 | 0.1% |
| S21-6 | 2785 | 80 | 0.5% | 0.2% | 7 | 0.3% |
| S21-7 | 2734 | 53 | 0.3% | 0.2% | 0 | 0.0% |
| S21-8 | 2204 | 48 | 0.4% | 0.1% | 0 | 0.0% |
| S21-9 | 3464 | 82 | 0.3% | 0.1% | 2 | 0.1% |
| S21-10 | 2595 | 73 | 0.2% | 0.3% | 6 | 0.2% |
| S21-11 | 2413 | 135 | 4.3% | 2.2% | 4 | 0.2% |
| S21-12 | 3467 | 82 | 0.3% | 0.2% | 5 | 0.1% |
| S21-13 | 2468 | 54 | 0.4% | 0.3% | 2 | 0.1% |
| S21-14 | 2147 | 50 | 0.2% | 0.1% | 4 | 0.2% |
| S21-15 | 2578 | 72 | 0.4% | 0.3% | 0 | 0.0% |
| S21-16 | 1523 | 39 | 0.3% | 0.3% | 0 | 0.0% |
| S21-17 | 2575 | 63 | 0.3% | 0.4% | 4 | 0.2% |
| S21-18 | 2313 | 57 | 0.2% | 0.2% | 0 | 0.0% |
| S21-19 | 3213 | 220 | 4.6% | 2.9% | 0 | 0.0% |
| S21-20 | 2667 | 66 | 0.3% | 0.3% | 3 | 0.1% |
| S21-21 | 3014 | 90 | 0.4% | 0.3% | 0 | 0.0% |
| S21-22 | 3102 | 71 | 0.4% | 0.1% | 0 | 0.0% |
| S21-23 | 2235 | 63 | 0.3% | 0.1% | 5 | 0.2% |
| S21-24 | 2669 | 56 | 0.1% | 0.2% | 4 | 0.1% |
| S21-25  (HT1-mCdHs-Cor1-1) | 2417 | 522 | 19.6% | 13.3% | 0 | 0.0% |
| S21-26 | 2372 | 66 | 0.5% | 0.2% | 0 | 0.0% |
| S21-27 | 2624 | 77 | 0.4% | 0.2% | 10 | 0.4% |
| S21-28 | 1934 | 75 | 0.7% | 0.5% | 0 | 0.0% |
| S21-29 | 2588 | 65 | 0.3% | 0.4% | 2 | 0.1% |
| S21-30 | 2399 | 63 | 0.2% | 0.2% | 0 | 0.0% |
| S21-31 | 2860 | 77 | 0.3% | 0.1% | 2 | 0.1% |
| S21-32 | 2157 | 53 | 0.5% | 0.2% | 2 | 0.1% |
| S21-33 | 2637 | 63 | 0.3% | 0.1% | 0 | 0.0% |
| S21-34 | 2608 | 60 | 0.3% | 0.2% | 2 | 0.1% |
| S21-35 | 2105 | 50 | 0.4% | 0.3% | 2 | 0.1% |
| S21-36 | 2642 | 71 | 0.5% | 0.2% | 5 | 0.2% |
| S21-37 | 2665 | 53 | 0.1% | 0.3% | 0 | 0.0% |
| S21-38 | 2330 | 48 | 0.2% | 0.2% | 4 | 0.2% |
| S21-39 | 2404 | 72 | 0.5% | 0.2% | 0 | 0.0% |
| S21-40 | 1247 | 26 | 0.3% | 0.5% | 2 | 0.2% |
| S21-41 | 2347 | 59 | 0.1% | 0.2% | 0 | 0.0% |
| S21-42 | 2306 | 57 | 0.3% | 0.3% | 4 | 0.2% |
| S21-43 | 2375 | 38 | 0.3% | 0.2% | 3 | 0.1% |
| S21-44 | 1956 | 191 | 7.2% | 3.3% | 0 | 0.0% |
| S21-45 | 2396 | 116 | 3.4% | 2.3% | 4 | 0.2% |
| S21-46 | 2227 | 56 | 0.4% | 0.1% | 2 | 0.1% |
| S21-47 | 2012 | 41 | 2.0% | 0.3% | 5 | 0.2% |
| S21-48 | 2608 | 67 | 0.3% | 0.1% | 2 | 0.1% |
| S21-49 | 2558 | 73 | 0.4% | 0.3% | 4 | 0.2% |
| S21-50 | 2527 | 44 | 0.2% | 0.2% | 0 | 0.0% |
| S21-51 | 2598 | 78 | 0.3% | 0.2% | 5 | 0.2% |
| S21-52 | 1506 | 31 | 0.0% | 0.3% | 0 | 0.0% |
| S21-53 | 2648 | 80 | 0.4% | 0.2% | 4 | 0.2% |
| S21-54 | 2482 | 63 | 0.3% | 0.2% | 0 | 0.0% |
| S21-55 | 2647 | 65 | 0.4% | 0.1% | 0 | 0.0% |
| S21-56 | 2241 | 63 | 0.5% | 0.2% | 0 | 0.0% |
| S21-57 | 3320 | 89 | 0.4% | 0.2% | 5 | 0.2% |
| S21-58 | 2964 | 82 | 0.4% | 0.2% | 0 | 0.0% |
| S21-59 | 2830 | 65 | 0.3% | 0.1% | 0 | 0.0% |
| S21-60 | 3539 | 80 | 0.4% | 0.2% | 0 | 0.0% |
| S21-61 | 3778 | 95 | 0.4% | 0.3% | 7 | 0.2% |
| S21-62 | 3229 | 105 | 0.4% | 0.2% | 10 | 0.3% |
| S21-63 | 3043 | 66 | 0.2% | 0.2% | 3 | 0.1% |
| S21-64 | 1744 | 39 | 0.3% | 0.1% | 2 | 0.1% |
| S21-65 | 2893 | 83 | 0.2% | 0.1% | 4 | 0.1% |
| S21-66 | 2694 | 62 | 0.2% | 0.0% | 0 | 0.0% |
| S21-67 | 3078 | 85 | 0.1% | 0.2% | 2 | 0.1% |
| S21-68 | 2568 | 66 | 0.3% | 0.2% | 4 | 0.2% |
| S21-69 | 3343 | 70 | 0.2% | 0.1% | 3 | 0.1% |
| S21-70 | 2955 | 62 | 0.2% | 0.2% | 0 | 0.0% |
| S21-71 | 2583 | 73 | 0.4% | 0.3% | 0 | 0.0% |
| S21-72 | 3237 | 72 | 0.2% | 0.2% | 3 | 0.1% |
| S21-73 | 2879 | 54 | 0.3% | 0.2% | 0 | 0.0% |
| S21-74 | 2698 | 65 | 0.2% | 0.1% | 0 | 0.0% |
| S21-75 | 2618 | 75 | 0.4% | 0.3% | 4 | 0.2% |
| S21-76 | 2033 | 56 | 0.3% | 0.4% | 3 | 0.1% |
| S21-77 | 2671 | 61 | 0.2% | 0.2% | 6 | 0.2% |
| S21-78 | 2502 | 57 | 0.3% | 0.2% | 2 | 0.1% |
| S21-79 | 2607 | 570 | 20.3% | 10.3% | 2 | 0.1% |
| S21-80 | 2427 | 51 | 0.3% | 0.0% | 6 | 0.2% |
| S21-81 | 3417 | 110 | 0.3% | 0.1% | 8 | 0.2% |
| S21-82 | 2713 | 51 | 0.1% | 0.3% | 0 | 0.0% |
| S21-83 | 2178 | 40 | 0.2% | 0.0% | 0 | 0.0% |
| S21-84 | 3356 | 73 | 0.5% | 0.3% | 3 | 0.1% |
| S21-85 | 2678 | 65 | 0.3% | 0.1% | 0 | 0.0% |
| S21-86 | 2356 | 58 | 0.6% | 0.3% | 0 | 0.0% |
| S21-87 | 2549 | 63 | 0.2% | 0.3% | 0 | 0.0% |
| S21-88 | 1588 | 36 | 0.1% | 0.2% | 0 | 0.0% |
| S21-89 | 3000 | 71 | 0.5% | 0.3% | 2 | 0.1% |
| S21-90 | 2640 | 61 | 0.3% | 0.2% | 2 | 0.1% |

**Table S2. Genomic sites captured by Digenome-seq**

|  | Chr : position | Cleavage score­­­ |
| --- | --- | --- |
| Control genomic DNA | chr19:60315442 | 6.37 |
|  | chr8:72112588 | 5.87 |
|  | chr3:28912995 | 5.46 |
|  | chr9:37000968 | 4.38 |
|  | chr18:60089364 | 4.26 |
|  | chr9:104123406 | 4.06 |
|  | chr19:60141777 | 4.01 |
|  | chr8:75136581 | 4 |
| ABE/EndoV-treated genomic DNA | chr3:117504849 | 6.01 |
|  | chr3:84545503 | 5.5 |
|  | chr9:74162358 | 5.45 |
|  | chr17:45840609 | 5.24 |
|  | chr19:37449202 | 5.02 |
|  | chr19:8955071 | 4.65 |
|  | chr19:60680279 | 4.62 |
|  | chr7:84595450 | 4.48 |
|  | chr7:84595450 | 4.47 |
|  | chr3:152059170 | 4.16 |
|  | chr7:16025764 | 4.12 |
|  | chr17:47517533 | 4.1 |

**­­­**

**Table S3. Primers for qRT-PCR (A) and high-throughput sequencing (B), related to STAR Methods.**

| **Gene** | **Primer** | **Sequence (5’ to 3’)** |
| --- | --- | --- |
| **A. Primers for** **qRT-PCR** | | |
| *Alb* | Fwd | GGCTACAGCGGAGCAACTGA |
|  | Rev | GCCTGAGAAGGTTGTGGTTGTG |
| *Sox9* | Fwd | TCCTAACGCCATCTTCAAGG |
|  | Rev | ACGTCTGTTTTGGGAGTGGT |
| *Epcam* | Fwd | TCGTGGTGGTGTTAGCAGTC |
|  | Rev | TCTGTGTATCTCACCCATCTCC |
| *Afp* | Fwd | CGTCCCTCCACCATTCTCTG |
|  | Rev | CGTGCTGCTCCTCTGTCATT |
| *Krt19* | Fwd | TTCCGGACCAAGTTTGAGAC |
|  | Rev | CCTCGTGGTTCTTCTTCAGG |
| *Itga6* | Fwd | GGATCATCCTCCTGGCTGT |
|  | Rev | TGTGGTAGGTGGCATCGTAA |
| *Cd44* | Fwd | GGCTTATCATCTTGGCATCC |
|  | Rev | CTGTTCCATTGCCACTGTTG |
| *Asgr1* | Fwd | CAGCTCTGTGAGGCCTTGGA |
|  | Rev | GGGCCCGTTCTGGTCAGTTA |
| *Hnf4α* | Fwd | ATCGRCAAGCCTCCCTCTGC |
|  | Rev | GACTGGTCCCTCGTGTCACATC |
| *Cyp1a2* | Fwd | AGGAGCTGGACACGGTGGTT |
|  | Rev | AGGTGTCCCTCGTTGTGCTG |
| *Cyp2c9* | Fwd | TGACTTGTTTGGAGCTGGGACAGA |
|  | Rev | ACAGCATCTGTGTAGGGCATGT |
| *Cftr* | Fwd | GGTCATAGAGCAGGGCAATG |
|  | Rev | TGCACTTCTTCCTCCGTCTC |
| *Ae2* | Fwd | GACTCCTTTCCCTGTGTGGA |
|  | Rev | GAAGCATCCGCTCTTTCTTG |
| *Aqpr1* | Fwd | CTGTGCGTTCTGGCTACCAC |
|  | Rev | GCACAGCAGAGCCAAATGAC |
| *Gabdh* | Fwd | CCAATGTGTCCGTCGTGGAT |
|  | Rev | TTGCTGTTGAAGTCGCAGGAG |
| **B. Primers for high-throughput sequencing** | | |
| *Fah* | 1st_Fwd | AGTAATGCCAGGTCCTCAGG |
|  | 1st_Rev | GTCAGCTCCATCCTTCCACT |
|  | 2nd_Fwd | ACACTCTTTCCCTACACGAC GCTCTTCCGATCT CTCCATGGCAGGCTTTCTTC |
|  | 2nd_Rev | GTGACTGGAGTTCAGACGTGT GCTCTTCCGATCT CCACACCCACAGAGTCAGAA |

**Table S4. Oligos for sgRNA plasmid cloning (A), pegRNA plasmid cloning (B), ngRNA plasmid cloning (C), and sgRNA *in vitro* transcription (D), related to STAR Methods.**

| **Gene** | **Primer** | **Sequence (5’ to 3’)** |
| --- | --- | --- |
| **A. Oligos for sgRNA cloning** | | |
| *Fah* | Spacer_up | CACCGACTGGAGCAGTAATGCCTGG |
|  | Spacer_down | AAACCCAGGCATTACTGCTCCAGTC |
| **B. Oligos for pegRNA cloning** | | |
| *Fah*_  peg1 | Spcaer_up | CACCGggatggtcctcatgaacgacGTTTT |
|  | Spcaer_down | CTCTAAAACgtcgttcatgaggaccatccC |
|  | 3’ext_PBS11nt_up | GTGCattacCgctccagtcgttcatgagga |
|  | 3’ext_PBS11nt_down | AAAAtcctcatgaacgactggagcGgtaat |
|  | 3’ext_PBS13nt_up | GTGCattacCgctccagtcgttcatgaggacc |
|  | 3’ext_PBS13nt_down | AAAAggtcctcatgaacgactggagcGgtaat |
|  | 3’ext_PBS15nt_up | GTGCattacCgctccagtcgttcatgaggaccat |
|  | 3’ext_PBS15nt_down | AAAAatggtcctcatgaacgactggagcGgtaat |
| *Fah*_  peg2 | Spcaer_up | CACCGtcagaggaagctgggccaccGTTTT |
|  | Spcaer_down | CTCTAAAACggtggcccagcttcctctgaC |
|  | 3’ext_PBS11nt_up | GTGCgcGgtaatgcctggtggcccagcttc |
|  | 3’ext_PBS11nt_down | AAAAgaagctgggccaccaggcattacCgc |
|  | 3’ext_PBS13nt_up | GTGCgcGgtaatgcctggtggcccagcttcct |
|  | 3’ext_PBS13nt_down | AAAAaggaagctgggccaccaggcattacCgc |
|  | 3’ext_PBS15nt_up | GTGCgcGgtaatgcctggtggcccagcttcctct |
|  | 3’ext_PBS15nt_down | AAAAagaggaagctgggccaccaggcattacCgc |
|  | Scaffold_up | AGAGCTAGAAATAGCAAGTTAAAATAAGGCTAGTCCGTTATCAACTTGAAAAAGTGGCACCGAGTCG |
|  | Scaffold_down | GCACCGACTCGGTGCCACTTTTTCAAGTTGATAACGGACTAGCCTTATTTTAACTTGCTATTTCTAG |
| **C. Oligos for ngRNA cloning** | | |
| *Fah*_  ng1 | Spcaer_up | CACCGTGGGCTTTGGAAATGGGGAT |
|  | Spcaer_down | AAACATCCCCATTTCCAAAGCCCAC |
| *Fah*_  ng1b | Spcaer_up | CACCGtacCgctccagtcgttcatg |
|  | Spcaer_down | AAACcatgaacgactggagcGgtaC |
| *Fah*_  ng2 | Spcaer_up | CACCGGCACACACAGGAGTTGGGTA |
|  | Spcaer_down | AAACTACCCAACTCCTGTGTGTGCC­­­ |
| *Fah*_  ng2b | Spcaer_up | CACCGgtcctcatgaacgactggag |
|  | Spcaer_down | AAACctccagtcgttcatgaggacC |
| **D. Oligos for sgRNA *in vitro* transcription** | | |
| *­Fah* | Fwd | GAAATTAATACGACTCACTATAGACTGGAGCAGTAATGCCTGGGTTTAAGAGCTATGCTGGAAAC |
|  | Rev | AAAAAAGCACCGACTCGGTGCCACTTTTTCAAGTTGATAACGGACTAGCCTTATTTAAACTTGCTATGCTGTTTCCAGCATAGCTCTTAAAC |
